# The chronology of developing cells: are epigenomic and transcriptomic oscillations linked to their linear trajectories?

**DOI:** 10.64898/2026.04.27.720664

**Authors:** Matthew Carlucci, Edward S. Oh, Turner Silverthorne, Adam R. Stinchcombe, Martynas Želnys, Artūras Petronis

## Abstract

**Background:** During development, the specification of individual cell types requires orchestrated shifts in the activity of thousands of genes, each following precise and coordinated trajectories. The ability of this process to succeed with remarkable reliability, despite its immense complexity, suggests that at least some underlying principles of development are fundamentally simple. Building on recent findings from epigenetic aging research, we hypothesize that linear trajectories in developing cells are influenced by concurrent oscillatory dynamics, which may help ensure synchrony and robustness.

**Results:** Supporting this model, we demonstrate an association between oscillatory and linear dynamics in cytosine modifications in mouse intestinal organoids, as well as in the transcriptomes of *C. elegans*. Furthermore, we show that transcriptomes of single cells exhibited developmental chrono-heterogeneity, enabling reconstruction of oscillatory cycles which also correlate with linear changes.

**Conclusions:** Oscillation-mediated linear dynamics may represent an evolutionary invention for encoding molecular time and orchestrating developmental processes.

## Background

Development and cell differentiation have been a focus of extensive research since the emergence of molecular biology. Half a million articles on this topic have been published over the last three decades, however, it remains a profound mystery how a fertilized oocyte gives rise to various tissues in a multicellular organism with such a remarkable success rate. Depending on the type of cell being formed, distinct combinations of thousands of genes must undergo precise, directional changes in a highly coordinated manner. Given the immense complexity involved in cell differentiation, *‘‘the amazing thing about animal development is not that it sometimes goes wrong, but that it ever succeeds’’* [1]. This precision, simplicity, and efficacy can be attributed to millennia of evolutionary refinement and optimization of developmental programs – metaphorically illustrated by C. H. Waddington’s epigenetic landscape, where the surface guides rolling marbles to determine their cellular fate (Additional file 2: Vid. S1). What are the molecular factors and mechanisms underlying the ridges, valleys, and gravity moving the marbles which assure the flawless and highly synchronized changes in the epigenomes of differentiating cells? We recently introduced the term ‘chrono-epigenetics’ as an umbrella concept encompassing the time-dependent dynamics of epigenetic processes [2]. New temporal connections emerged from the observation of diurnal oscillations in cytosine modification densities; such cytosines also exhibited age-dependent linear modification changes [3,4]. Quantitatively, the parameters of epigenetic oscillations predicted how they were aging: acrophases (the time of a cycle’s peak) indicated the direction (gain or loss of modification) of epigenetic aging, while oscillation amplitudes correlated with the magnitude of linear changes [3,4]. These findings support the idea that cyclical and linear processes in the epigenome are interconnected (Fig. 1A). Given that aging and development are facets of the same biological process [5], principles of the former may be applicable to the latter. Instead of convergence, or regression to the mean, in the aging cells, epigenomes may diverge in the developing cells (Fig. 1B). Directly relevant is the observation that the epigenetic clocks, which were originally developed to measure chronological and biological age in humans and other animals, in fact, start ticking quite accurately from early stages of embryogenesis [6,7]. Oscillations may be a shared feature of both development and aging, with the primary formal distinction between the two processes lying in which specific genes, transcripts, and proteins oscillate – and in the characteristics of those oscillations, such as their phases and amplitudes (Fig. 1A versus B).

**Fig 1.**
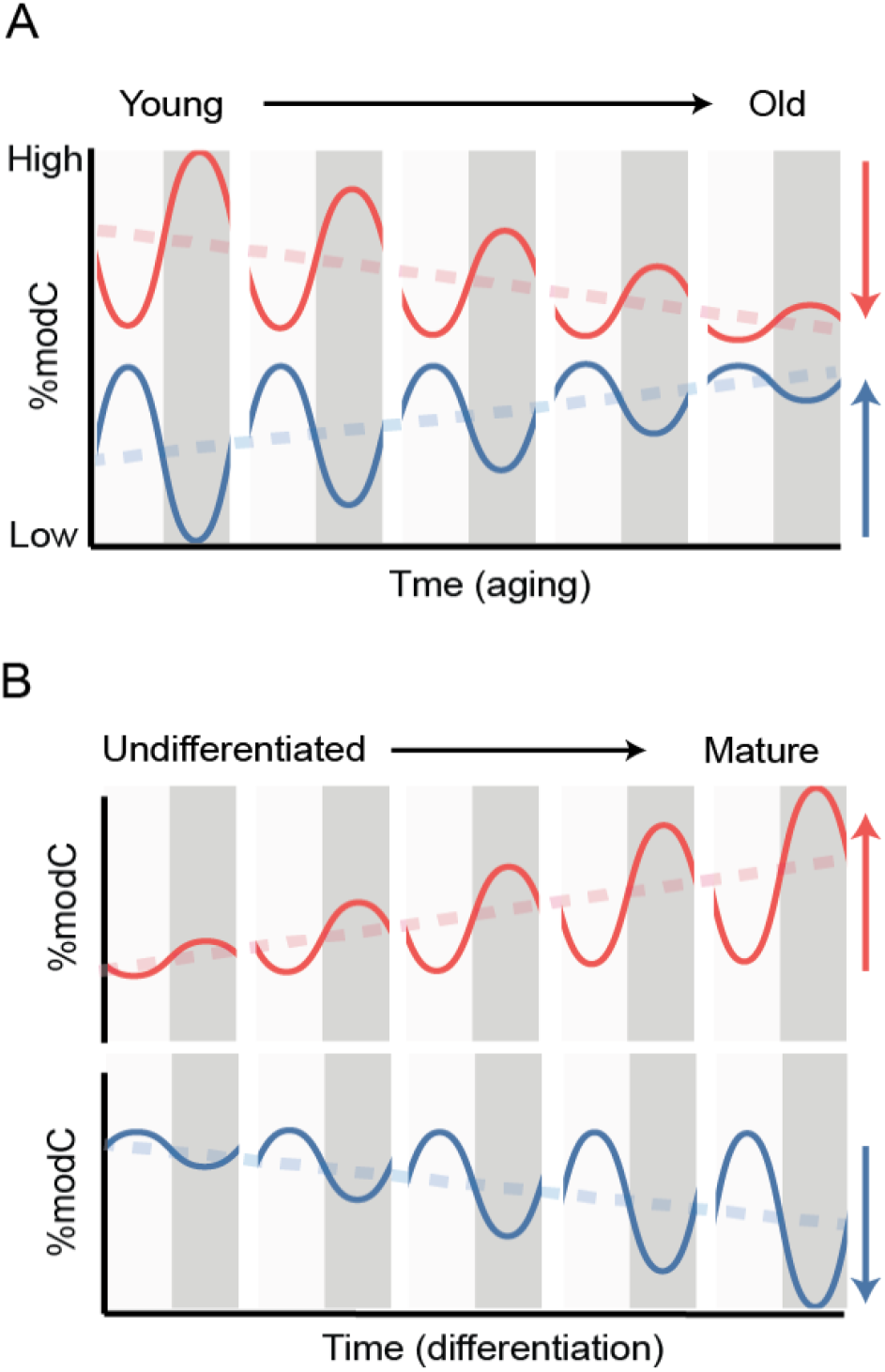
Hypothetical oscillatory-linear dynamics in the development-aging continuum. **A** In aging, phases of oscillations account for the direction of age-dependent epigenomic changes and convergence of polarized cytosine modification densities, also known as regression to the mean. **B** In development and cell differentiation, epigenomic divergence could arise from the same oscillatory-linear principle but with different loci, or different oscillation phases and amplitudes.

Here, we propose the foundational basis for the <u>chr</u>ono-<u>e</u>pigenetic model <u>o</u>f <u>d</u>evelopment (CHREOD), positing a principle for orchestrating directional, synchronized, and pre-programmed molecular changes during cell differentiation. It is worth noting that the term ‘chreod’ (or ‘creode’), which conveys a similar concept but has a vastly different etymology, was introduced into developmental biology seven decades ago [8]. Waddington coined this neologism from the Greek roots for ‘necessary’ and ‘path’ to emphasize the deterministic pathway that guides differentiating cells. In an adaptation to Waddington’s chreod, our CHREOD suggests that the marbles do not travel in straight lines but swing rhythmically, with periodicities and velocities aligned with the structure of the ridges, guiding them along the ‘necessary paths’ (Additional file 2: Vid. S2).

Although initial insights for CHREOD came from DNA modification studies, the principles discussed below are not limited to canonical epigenetic marks, i.e. modifications of DNA and histones. Indeed, the molecular layers and elements of cell organization such as RNA, histone modifications, transcription factors, proteins, metabolites, and chromatin conformation are inseparably interconnected and regulated by one another. Any layer or element of the oscillating ‘multi-geared’ network provides insights into cellular dynamics during cell development. This perspective is in line with the historical definitions of epigenetics which include *all* mechanisms governing cell differentiation and establishing new cellular identities [9]. Therefore, in addition to DNA and histone modifications, CHREOD encompasses all other intracellular components, most notably, RNA and proteins.

Several types of molecular oscillators have been detected in embryonic development [10,11]. The classical example is the segmentation clock, where the period of *Hes/Her* mRNA oscillations dictate temporal cues for the segmentation of somites in the presomitic mesoderm. Two-three hour oscillations of modified cytosines [12] and nascent RNA [13] were detected in stem cells. In neural progenitor cells, Hes1, Ascl1, and Olig2 oscillate, change linearly, and contribute to differentiation into astrocytes, neurons, and oligodendrocytes [14,15]. Linear trends among oscillating transcripts were observed in several other studies [14,16,17] (Additional file 1: Fig. S1), however, their interrelationships have never been systematically explored.

In this study, we utilized temporal epigenomic and transcriptomic approaches to test the core element of CHREOD — the relationship between molecular oscillations and linear trajectories during development and cell differentiation. Using mouse intestinal organoids, we investigated how epigenomic oscillations are related to development and transitions in cell identity, from intestinal stem cells to mature enterocytes. We also investigated cyclical-linear dynamics between the ultradian transcriptomic oscillations and its various cell lineage outcomes from embryogenesis to the young adult stage of the nematode *C. elegans*. Finally, we employed novel methodologies to detect oscillatory-linear links in a single timepoint single-cell transcriptomic dataset, which is not feasible with conventional approaches.

## Results

### Oscillating-linear associations in the epigenomes of differentiating mouse enterocytes

To test the hypothesis that oscillations in the epigenome are related to linear changes, we utilized intestinal organoids generated from mouse duodenum [18]. Circadian rhythms of individual mouse organoids (n = 45) were confirmed using a clock fluorescence mPeriod1-Venus reporter over 144 h (Fig. 2A-C). 24 h oscillations were the dominant periodicity across most organoids; 33 out of 45 organoids’ highest R^2^ was for 24 h periodicity, and 42/45 organoids were between 22 to 26 h (Fig. 2B). The acrophases of mPer1-Venus were synchronized around ZT12 (min-max = 10.3–14.9) (Fig. 2C). *Per1, Per2*, and *Arntl* mRNAs were also confirmed for circadian oscillations by qPCR (Fig. 2D).

**Fig. 2.**
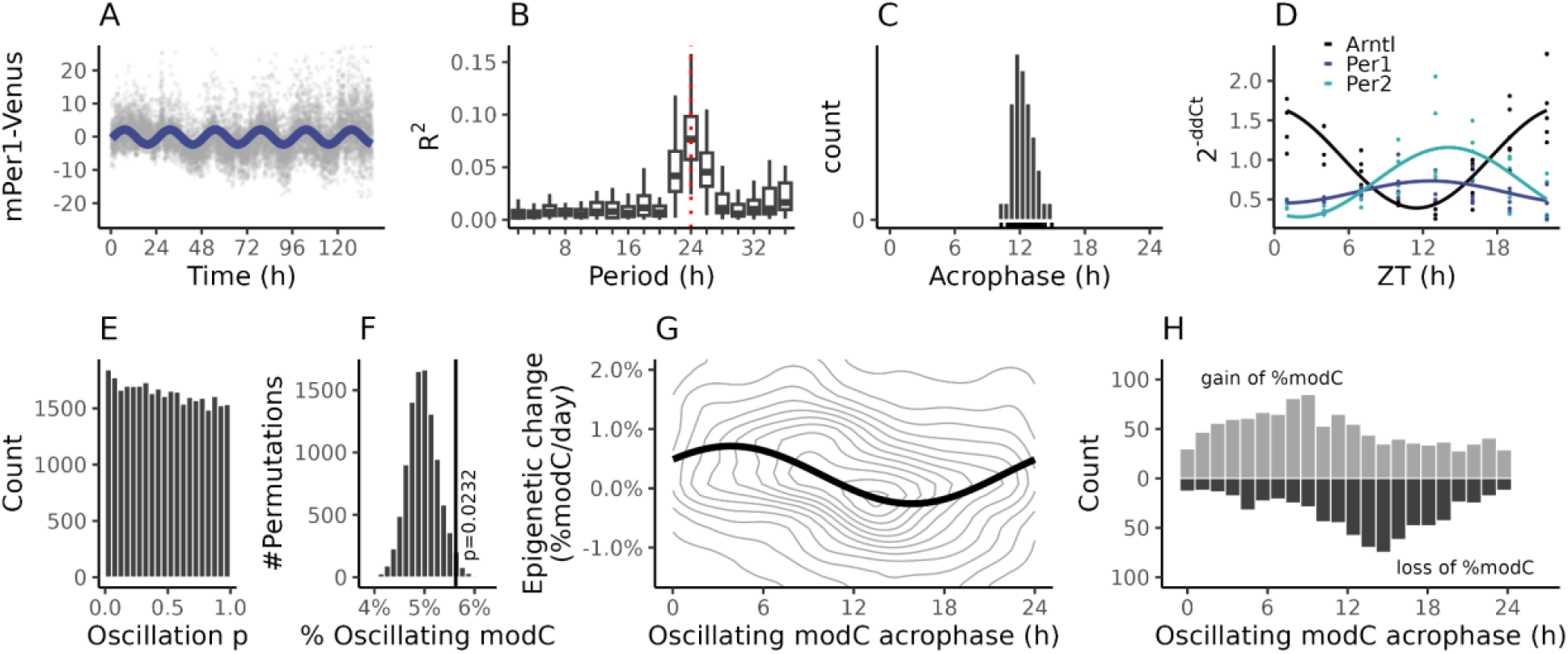
Circadian modCs are associated with linear epigenetic changes. **A-D** Circadian gene expression in mouse intestinal organoids. **A** mPer1-Venus luminescence intensity recordings of intestinal organoids (n = 45 organoids). The cosinor curve (blue) was estimated across all organoids with a period of 24 h. **B** Periodogram of mPer1-Venus signal across individual organoids (n = 45). **C** mPer1-Venus acrophase distribution for 24 h periodicity across individual organoids. **D** 24 h oscillations, with cosinor fits, of core circadian gene (*Arntl, Per1, Per2*) transcripts using qPCR with mouse intestinal epithelium marker *Villin-1* as endogenous control. **E-F** Circadian oscillation results of modC interrogated by targeted bisulfite sequencing. **E** Cosinor *p*-value histogram of individual cytosines. **F** Proportion of cytosines with oscillating modC by harmonic regression fits across all cytosines in each of 10,000 permutations. The vertical line indicates the observed proportion (5.6%) with the *p*-value indicating the proportion of permuted tests that exceeded the observed. **G** Association between acrophases of oscillating modCs (X-axis) and estimates of epigenetic linear change (Y-axis). Contours (grey) were rendered using 2D kernel density estimation. The cosinor curve (black) was estimated by regression, using the sine and cosine of the acrophase as predictors of linearity. Plot is truncated at -1% and 2% from the full data range of - 11.1% to 7.3%. **H** Bidirectional histogram of oscillating modC acrophases demarcated by those that were gaining (top) or losing (bottom) modifications with time.

In order to obtain circadian sampling, individual culture wells of mouse intestinal organoids were harvested every 4 h for 32 h (n = 1–3 biological replicates per time point after quality control, total n = 21, see Methods). Extracted DNA was subjected to targeted bisulfite sequencing [19] in technical triplicate at an average samplewise depth of 521 ± 90 (mean ± SD) reads per cytosine to obtain cytosine modification (modC) densities (Additional file 3: Table S1). Using a cosinor model [20] with linear and batch covariates (see Methods), we detected 24 h oscillations in the modification densities of 1,843 out of 32,767 cytosines (5.6%; cosinor *p* < 0.05) (Fig. 2E; Additional file 1: Fig. S2A). Individual cytosines did not survive correction for multiple testing, so we performed a permutation test and found global 24 h oscillation significance (permutation *p* = 0.023, see Methods) (Fig. 2F). The linear component of these oscillating modCs was used as an estimate of modification change that occurred during intestinal epithelial development over 32 h in culture (Additional file 1: Fig. S2B; Additional file 1: Fig. S3).

To test whether phases of oscillating modCs connect to linear epigenetic changes, we correlated the acrophase of oscillations against the linear rate using a second cosinor model. We detected a small but significant association (R^2^ = 0.06, cosinor *p* = 1.1×10^−23^) (Fig. 2G) indicating that oscillating modCs with acrophases at ZT4 ± 6 and the antiphasic 16 ± 6 were more likely to increase and decrease in their modification densities, respectively (Fig. 2H).

### Putative linear epigenomic effects arising from the synergy of circadian and cell cycles

We explored whether the known molecular synchronization between the circadian clock and the cell cycle [21,22] can explain how epigenetic oscillations facilitate linear epigenomic changes during differentiation. Our model is based on two assumptions. First, epigenetic oscillations pause during cell division, e.g. when chromatin condenses into visible chromosomes in the G2/M phase. Second, based on the dominating 24 h periodicity (Fig. 2A-C), oscillations in the daughter cells resume in circadian synchrony with the rest of the organoid. As a result, a linear effect emerges from changes in the MESOR (Midline Estimating Statistic of Rhythm) (Fig. 3A), and this change depends on the oscillator’s parameters as well as the timing of the pause window (see Methods). That is, oscillations pausing during the high velocity ascending or descending phases should decrease or increase their MESOR, respectively (Fig. 3B). In contrast, oscillations pausing at a lower velocity phase, i.e. close to their acrophase or nadir, should accumulate less, or no linear change (Fig. 3B).

**Fig. 3.**
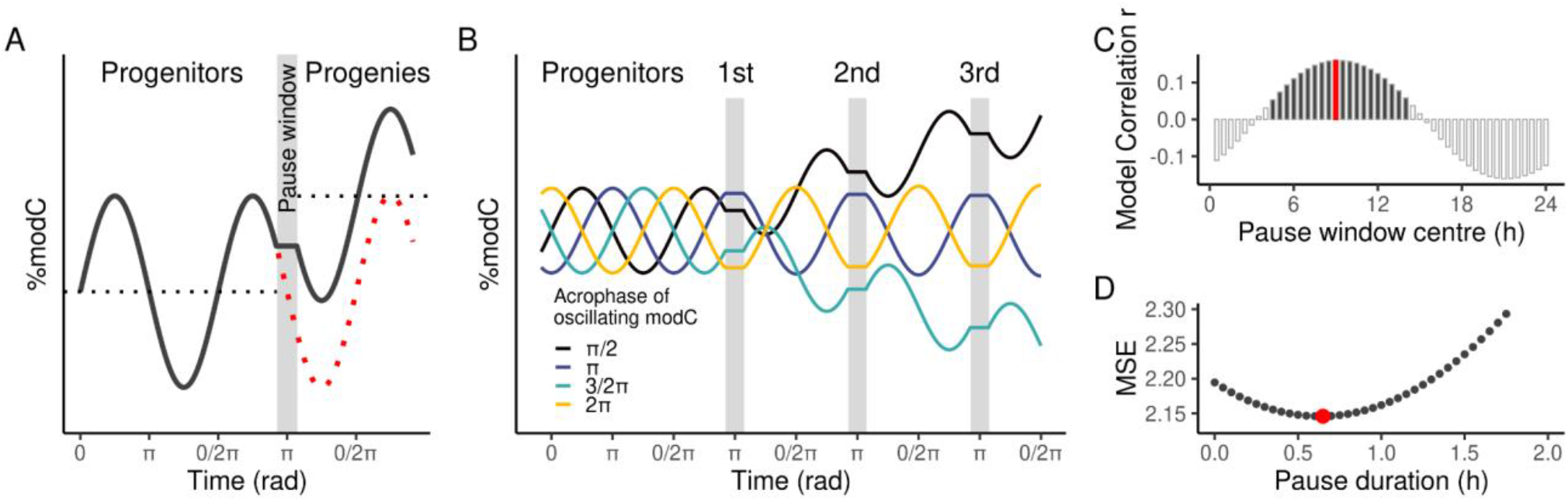
CHREOD dynamics explained by a circadian and cell cycle synchronization model. **A** Oscillating modCs in the progenitor cells may pause (grey box) during cell division resulting in a diversion from its normal trajectory (red) to a trajectory (black) with a shifted MESOR (black dotted line). A hypothetical pause window during one eighth of the period (centred at π and spanning π/4) generates an increasing MESOR in an oscillation with acrophase of π/2. **B** Same as **A**, but for 4 types of modC oscillators across 3 cell cycles. Oscillating modCs pausing during a high velocity phase (black and green) display MESOR changes. The MESORs of oscillating modCs pausing at their acrophase (blue) or nadir (yellow) do not change. **C** Pearson’s correlation *r* of the model predicted linearity with observed linearity (Y-axis) for all possible window centres (X-axis). The best model (red bar) uses a window centre of ZT9, yet other window centres were significant (*p*<0.05; dark grey bars). This correlation was unaffected by pause duration (Additional file 1: Fig. S4). **D** Tested pause durations (X-axis) with a window centre of ZT9 and their corresponding mean squared error of the model (Y-axis). The pause duration with the best mean squared error (MSE = 0.021%) is shown in red.

Since the pause window centre and duration were unknown, we investigated which windows could explain the observed modC linearity based on the observed oscillating modC amplitudes and acrophases. All windows with a centre between ZT5 and ZT14 generated linearity predictions that significantly correlated with the data (Pearson’s *r* = 0.058–0.16; *p* = 2.2 × 10^−12^–6.2 × 10^−3^), with the strongest one centred at ZT9 (Fig. 3C). The ZT9 model was found to have the least error when the interval spanned 0.65 h (Fig. 3D). In line with the primary oscillation-linear analysis (Fig. 2G and H), a ZT9 pause model yielded oscillating modCs with acrophases at ZT3 and ZT15, inducing the most pronounced linear effects in the positive and negative directions, respectively. These findings indicate that under such a model, even short pauses in epigenetic oscillations can drive linear changes of modCs, highlighting a potential mechanism by which circadian and cell cycle interactions shape the differentiating epigenetic landscape.

### Oscillating-linear associations in the transcriptome of developing *C. elegans*

To determine whether CHREOD generalizes to other organisms, we investigated the oscillatory-linear links in the developing nematode *C. elegans*. While the classical epigenetic factor, 5-methylcytosine, is lacking in *C. elegans*, transcriptomes start oscillating in the embryo and continue throughout larval development [23]. We utilized three transcriptomic datasets — larval bulk RNA-seq [23] as well as larval and adult single-cell RNA-seq [24,25] — which allowed us to test CHREOD at both the whole-organism and tissue-specific levels (Additional file 1: Fig. S5).

For the whole-worm analysis of oscillating-linear associations, oscillatory acrophases were obtained from the synchronized larvae bulk RNA-seq dataset. The hourly sampling revealed that 3,739 of 15,371 (24%) mRNA transcripts oscillated with a 7 h periodicity (using the original study’s criteria: cosinor *p* < 0.01 and fold-change amplitude > 1.4) during larval stages L2 to L3 [23]. Linear mRNA changes from embryogenesis to young adulthood were estimated from the adult scRNA-seq dataset [25]. For the start point, we used oocytes which were collected in large numbers (n = 4,447 cells) and whose transcriptomes are similar to early embryos [26] (Additional file 1: Fig. S5B). The transcriptomes of all somatic cells in the adult worm at 55 h post-hatching were used as the endpoint (n = 94,219 cells), with their differences relative to oocytes estimating the linear changes (change = RNA_soma_ − RNA_oocyte_) (Additional file 1: Fig. S5C). Consistently with CHREOD predictions, acrophases of the 3,739 oscillating *C. elegans* mRNAs were associated with their linear changes (cosinor *p* = 5.3 × 10^−40^) (Fig. 4A). Antiphasic oscillating mRNAs, reaching acrophases at 4.7 h and 1.2 h of the cycle, were associated with the most upregulation and downregulation, respectively. However, the proportion of the linear change explained by oscillation phases was quite small (R^2^ = 0.048) which may have been due to the averaging of tissue-specific oscillatory and linear parameters.

**Fig. 4.**
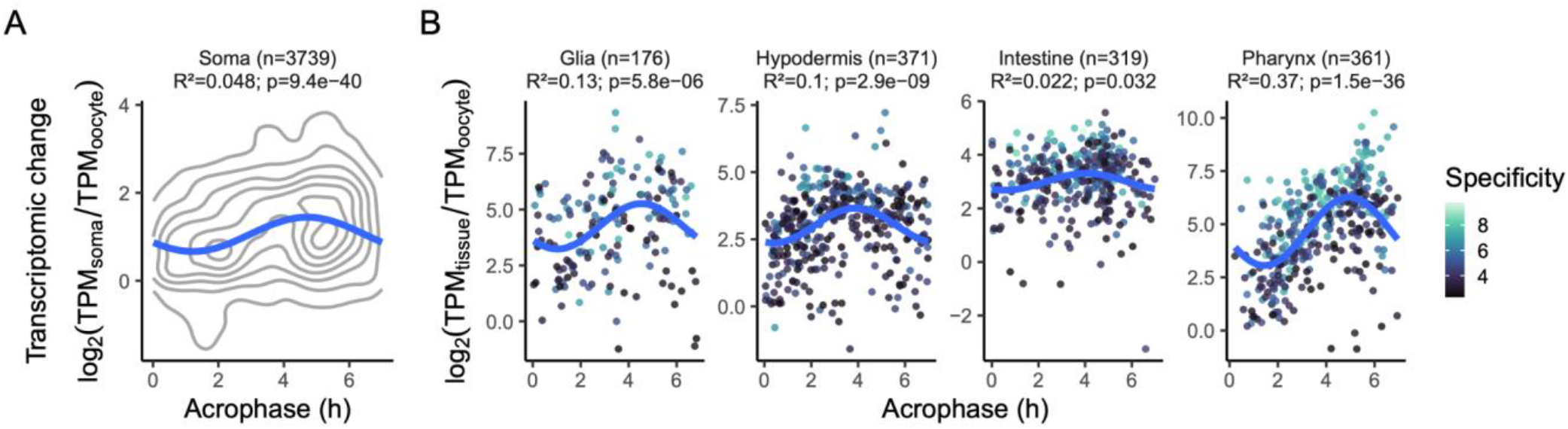
The strength of the oscillatory-linear association increases from the whole-worm to the tissue-specific context. **A** Whole-worm association between acrophases of oscillating mRNAs with a 7 h period (X-axis) and estimates of transcriptomic linear change of somatic cells (Y-axis). Linearity was the difference in logarithm (base 2) transcripts per million between oocytes and all somatic cells. Contours (grey) were rendered using 2D kernel density estimation. The cosinor curve (blue) was estimated by regression, using the sine and cosine of the acrophase as predictors of linearity. n - number of oscillating transcripts. **B** The same association as in **A** but for tissue-specific genes; with separate panels for each of the four tissues: glia, hypodermis, intestine, and pharynx. Linearity here was the difference in logarithm transcripts per million between the tissue and oocytes. Color of each point indicates the degree of specificity of the tissue-specific genes as the logarithm (base 2) of the transcript ratio relative to the second highest cell type obtained from the original study [24]. Tissue specificity was added in the association test as a covariate, and R^2^ are partial R^2^ after accounting for the variance explained by specificity. n - number of oscillating transcripts.

To test whether individual tissues yield a higher R^2^, we selected hypodermis, pharynx, glia, and intestine which contained the highest number of oscillating mRNAs (n = 176–371) that were tissue-specific in the larval scRNA-seq dataset [24]. Linearity was calculated within the tissue-specific genes of each tissue (n = 5,534–14,985 cells) using the same contrast to oocytes as before (Additional file 1: Fig. S5). All tissues demonstrated a significant association between phase and the degree of linear change (cosinor *p* = 1.5 × 10^−36^–0.032) (Fig. 4B). Consistent with the prediction regarding deflated R^2^ in the whole worm, the proportion of linear dynamics explained by oscillation phases in the tissues increased substantially (R^2^_partial_ = 0.022–0.37; average R^2^_partial_ = 0.16).

### Oscillating-linear associations in chrono-heterogeneous single cell transcriptomic datasets

The vast majority of developmental studies capture snapshots at only limited stages, making it difficult to reliably detect oscillations. Single-cell sequencing technologies, however, reveal that cells collected at a single time point can exhibit developmental time variability, and this ‘chrono-heterogeneity’ may be sufficient to reconstruct full oscillatory cycles [27–29]. We established such approaches for testing CHREOD by applying them to a scRNA-seq dataset sampled at the larval L2 stage [24]. Utilizing the existing cell annotations [30], we investigated the 33 most abundant cell types (n > 200 cells in each) in the dataset. This enabled testing of the previous whole-worm, and tissue-level CHREOD associations at the individual cell type level (Additional file 1: Fig. S5).

Cycle reconstruction utilizes the circular signature that emerges from the correlation of two oscillators phase-shifted by a quarter cycle (Fig. 5A and B). We started by performing PCA to project the transcriptome of each cell onto a two-dimensional PC1-PC2 space for each cell type (Additional file 1: Fig. S6A). A linear regression-based ellipse fit (see Methods) was then performed to determine whether the cell arrangement is consistent with two phase-shifted oscillatory signals (permutation *p* < 0.05) (Fig. 5C). Additionally, an ellipse fit threshold (R^2^ > 0.2) was used to determine which cell types had accurate phase estimates suitable for further analysis (Additional file 1: Fig. S6B). For example, anterior arcade cells had a strong ellipse fit suggesting a cyclical arrangement (permutation *p* < 0.001; R^2^ = 0.44). Anterior body wall muscle cells were also significant for a circular topology (permutation *p* < 0.05), however, its ellipse fit quality was too low (R^2^ = 0.0057) for accurate phase inference. Other cell types such as the posterior body wall muscle cells, despite their large sample size (n = 1,088), provided no evidence for PC1-PC2 based oscillations (permutation *p* = 0.48). Out of the selected 33 cell types analyzed in the dataset, 5 cell types, namely seam, posterior/anterior arcade, hypodermal, and excretory duct cells, had high quality ellipse fits (permutation *p* < 0.001; R^2^ = 0.27–0.47) (Fig. 5D) and the cell phases along each ellipse were used for further analysis.

**Fig. 5.**
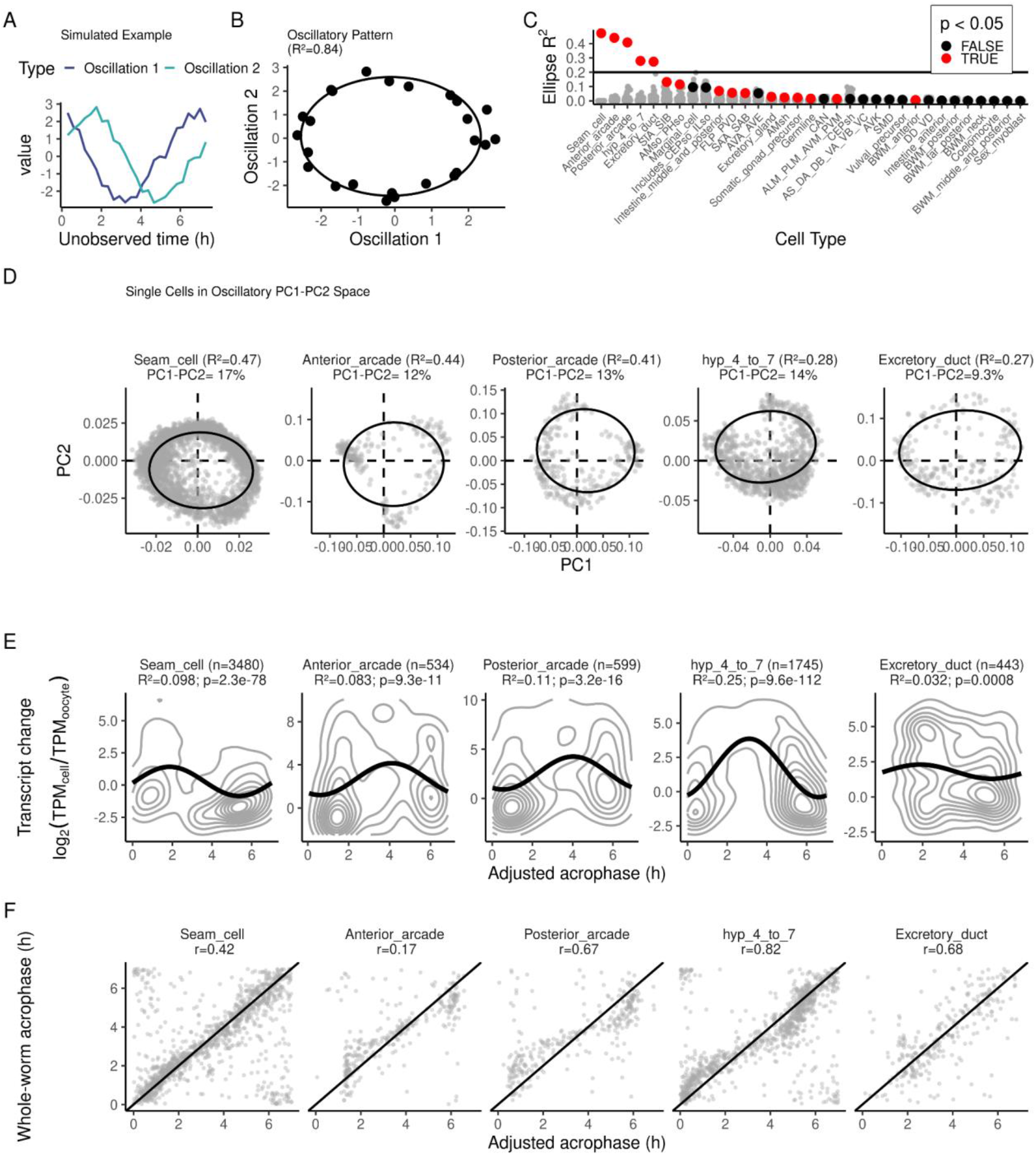
Chrono-heterogeneity in single cell transcriptomes reveals oscillations linked to linear trends. **A** Two simulated oscillatory signals (Y-axis) with a phase shift of a quarter of a cycle shown over an unobserved time variable (X-axis). **B** Correlation of signal 1 and 2 from **A** with an ellipse fit shown (line) and ellipse R^2^ reported. **C** Observed ellipse R^2^ (Y-axis; red/black dots) versus 100 representative null permutations (grey dots) for all investigated cell types (X-axis). Red dots were significant (permutation *p* < 0.05) for the ellipse permutation test versus 10,000 null permutations. Cell types are ordered by observed R^2^. **D** PC1 (X-axis) and PC2 (Y-axis) for cell types where PC1-PC2 displayed a significant ellipse fit; R^2^ of ellipse and combined variance explained by PC1-PC2 (compared to the total of the first 50 PCs). **E** Cell-level association between adjusted acrophases of oscillating mRNAs with a 7 h period (X-axis) and estimates of transcriptomic linear change of the corresponding adult cell type (Y-axis). Linearity was the difference in logarithm (base 2) transcripts per million between oocytes and the adult cell population. Contours (grey) were rendered using 2D kernel density estimation. The cosinor curves (black) were estimated by regression, using the sine and cosine of the acrophase as predictors of linearity. n - number of oscillating transcripts. Prior to acrophase adjustments, acrophases were reflected and/or shifted, but association strength was unchanged (Additional file 1: Fig. S8A). **F** Adjusted acrophases (X-axis) correlated with whole-worm acrophases (Y-axis). Correlation value (r) from a circular correlation test [31,32] is shown in each panel and all were *p* < 6.8 × 10^−4^. Single-cell acrophases were adjusted by a reflection and/or shift to minimize the phase difference relative to the whole-worm (see Methods), but correlation strength (r) and significance was unchanged compared to the unadjusted data (Additional file 1: Fig. S8B). **D-E** Panels are ordered according to the ellipse R^2^ of each cell type.

By applying cosinor modeling to the scRNA-seq transcriptomes of the five reconstructed cell types, we identified 4,459 oscillating transcripts (cosinor FDR *q* < 0.05), of which 1,755 corresponded to whole-worm oscillating transcripts while 2,704 were detected as putative new oscillators (Additional file 1: Fig. S7). Linear transcriptomic changes were estimated as in the previous analyses, i.e. oocytes were used as the starting point and the corresponding cell types in the adult worm served as the endpoint (n = 820–3,076) (Additional file 1: Fig. S5). We detected significant associations of cell type level oscillations with linearity in all 5 cell types (cosinor *p* = 9.6 × 10^−112^–8.0 × 10^−4^; R^2^ = 0.032–0.25) (Fig. 5E; Additional file 1: Fig. S8A).

Ellipse fits traced each cell type’s oscillatory cycle, yet without a reference, the absolute time for a given phase, and the direction of the cycle, remained unknown. To obtain these time and direction references on the cycle, we compared gene-wise acrophase estimates to the whole-worm oscillatory data and assumed that oscillations on average would be aligned with the whole-worm oscillations. Oscillating transcripts in all 5 cell types significantly overlapped with the whole-worm oscillations (all Fisher’s exact test *p* < 10^−60^; OR = 2.1–8.3). Among these overlapping oscillating transcripts, the acrophases exhibited a strong correlation (|circular *r*| = 0.17–0.82; all circular correlation *p* < 6.8 × 10^−4^), supporting the cycle reconstruction accuracy and the assumed average coherence with the whole worm. Then, to resolve the time and direction along each cycle, cell phases were globally phase-shifted and/or reflected within each cell-type in order to best match the whole-worm acrophases (Fig. 5F; Additional file 1: Fig. S8B).

With each cell type aligned to the same whole-worm reference, we detected that 31-88% of oscillating transcripts exhibited statistically significant differences in oscillations between pairs of cell types (pairwise differential oscillation FDR *q* < 0.05). Even similar cell types such as anterior and posterior arcade cells or hypodermal and seam cells had small average acrophase differences of 0.5 h and 0.8 h, respectively (Additional file 1: Fig. S9). Across these types, however, average pairwise acrophase differences were much larger and reached 1.2–1.4 h. This difference, which is not too far from a total phase mismatch (i.e. 1.75 h for a 7 h cycle), may have hindered oscillation detection in the whole-worm analysis. Therefore, using a single scRNA-seq sample, we uncovered evidence of oscillatory heterogeneity across tissues during *C. elegans* development which was not apparent in whole-worm analysis but is critical for detection of cell type-specific CHREOD effects.

## Discussion

The question we posed — whether shared molecular principles underlie development and cell differentiation — is not new. Over 50 years ago, the *C. elegans* project was initiated to uncover the logic by which gene networks regulate the development of diverse cell types in multicellular organisms [33]. A decade later, it was concluded “*there is not a simple set of obvious rules that governs development*” [34]. Since then, extensive experimental data on *C. elegans* and many other organisms has been accumulated, providing renewed motivation in the search for fundamental principles of development. However, it is important to recognize that even large volumes of data may not, on their own, be sufficient to unravel the complexity of developmental programs. New conceptual frameworks may be essential to fully understand how cells orchestrate their differentiation and fate decisions [35].

For any meaningful generalization, it is essential to choose the appropriate level of coarseness that avoids getting lost in details while not letting abstraction overshadow the sought core phenomena. We conceptualized development as numerous directional changes in the levels, amounts or densities of epigenomic, transcriptomic, proteomic, and other factors. The challenge, as we see it, lies in determining which factors will increase or decrease and the timeframe within which these synchronized changes will occur. The CHREOD model proposes that oscillations orchestrate these linear processes, ensuring synchronized and timely progression of the multilayered cellular machinery.

Our study of mouse intestinal organoids and *C. elegans* showed that epigenomic and transcriptomic trajectories in developing cells are indeed coupled with oscillatory dynamics. ‘Zigzagging’, instead of straightforward moving from start to end, may provide cells with a framework for encoding and interpreting temporal information [36]. Notably, oscillations, with their interwoven feedback loops, contribute to both the robustness of developmental processes and their ability to adapt to perturbations [37]. Any delay or acceleration of one oscillating element can be corrected by the rest of the multigear system, ensuring synchronized progression toward multiple molecular destinations. The presence of antiphasic communities observed in both epigenomes and transcriptomes may further enhance robustness [38,39]. Feedback mechanisms that control and regulate oscillations are likely to engage multiple ‘omics’ layers. Transcription-translation feedback loops thoroughly investigated in circadian biology [40] would not be possible without epigenetic, metabolic, and other diverse components. While initial insights were drawn from studies on DNA modifications, we suggest that the principles of CHREOD may extend across the other levels of cellular biology ‘above’ DNA: encompassing RNA, transcription factors, DNA-binding proteins, metabolites, and other components of the multigear oscillatory system in the differentiating cell.

Mechanisms generating linearity may be numerous and interrelated with one another. We proposed that the cell cycle, when coupled with circadian epigenomic oscillations, can drive directional changes across progeny. This resulted in a simple model based on iterative directional shifts in the MESOR of progeny cells that emerge from pauses in epigenetic oscillations. Pause windows between ZT5–ZT14 generated predictions consistent with linear modC changes in intestinal organoids. Relatedly, a simple phase arrest, which was a focus of the *C. elegans* oscillator study [23], can generate linear effects without any cell cycle context, e.g. an oscillator arresting at acrophase or nadir would produce higher, or lower linearity, respectively. Such a phase arrest-based mechanism exhibits the restricted feature that linear effects are no larger than the amplitude of oscillation, i.e. they are within the limit cycle in phase space. Two other mechanisms of linearity are worth mentioning. If co-existing but functionally opposing factors—such as DNA methylation and demethylation enzymes [41,42] — are imbalanced, the dominant factor can cause a gradual shift in the MESOR. Linear effects can also be generated by kinetic differences among components of the transcription–translation machinery. For instance, oscillatory transcription factor binding would drive cyclical production of target RNA transcripts, but their degradation may be slower than that of the transcription factor itself, leading to accumulation of RNA—and, subsequently, its protein product [43,44].

While the reported results highlight a prevalent oscillatory-linear association, each of our datasets also contained a subset of oscillators exhibiting the opposite direction of linearity. It remains unclear whether these deviations stem from experimental limitations, the coexistence of two oscillation-linear dyads, or other yet unknown factors. Among the experimental limitations, neither the linear nor the oscillatory measurements were fully reliable for accurate probing of the start, oscillatory, and end point cell populations. In *C. elegans* studies, the oocyte as a starting point and the adult stage as an endpoint were likely too early and too late, respectively, to capture exclusively oscillatory-linear dynamics. Oscillatory analyses of both worm and intestinal organoids represented mixed cell populations, a limitation mitigated by exploiting chrono-heterogeneity within individual cell types of *C. elegans* larvae. However, technical limitations of single-cell technologies and non-oscillatory influences to the cycle reconstruction inevitably affected the precision of the experiment, e.g. cycles were successfully reconstructed in only 5 out of 33 cell populations. Oscillations may exist in other cell types but could have been masked by undetected cell heterogeneity in the scRNA-seq annotation, low cell or gene counts, or more complex temporal signals than multiphasic oscillations. These issues underscore the need for concrete refinements in both experimental design and analytical methods.

The CHREOD framework and its associated mechanisms posit that oscillations drive linear effects. To prove their causative relationship, a series of experiments need to be performed, e.g. monitoring molecular oscillations throughout cell cycle phases, examining interactions among competing gene regulatory systems, and assessing distal effects of oscillators. Additional proofs of causality may involve experimenting with disturbed cycles of selected targets and finding out how it affected linear changes. Such experiments may help in understanding other important elements of CHREOD, namely the robustness and self-correcting nature of interconnected cycles. How would disturbed oscillators affect the rest of the oscillatory machinery, and *vice versa*, to what extent would the disruption be rectified?

The cause-and-effect relationship between oscillations and linear processes may not be unidirectional. While CHREOD is primarily focused on how oscillations orchestrate linear processes, the opposite direction of regulation is also possible. Increasing or decreasing MESORs may eventually change oscillation amplitudes and even quench oscillations completely. A prototype of such a mechanism can be found in differentiating neural stem cells where Neurog2 oscillations induce linear accumulation of Tbr2, which subsequently suppresses Hes1 — an oscillator antiphasic to Neurog2 [15].

Understanding the emergence and synchronization of oscillations was beyond the scope of this study. This is not a unique challenge, however, and valuable insights can be drawn from a wide range of biological systems such as networks of pacemaker cells in the heart, circadian pacemaker cells in the suprachiasmatic nucleus in the brain, metabolic synchrony in yeast cell suspensions, among numerous others [45]. In well-characterized developmental pathways, such as cAMP oscillations in *Dictyostelium* cells, mechanistic models [46] predict that both positive and negative feedback loops are required for the emergence of oscillations after stochastic cAMP signaling [47]. Larger oscillatory networks are often described by the Kuramoto model [45], in which individual oscillators evolve autonomously with a free-running period and interact via a sinusoidal coupling function. If the coupling is sufficiently strong to overcome heterogeneity among individual oscillators, the network can reach a synchronized state—an idea that may be applicable to developing cells [12]. Beyond the emergence of synchrony, additional features such as oscillator excitability [38] and community organization [39] can be added to produce the antiphasic states and synchrony necessary for CHREOD dynamics.

Our experimental findings and theoretical insights suggest that the foundational CHREOD framework may be reducible to self-regulating oscillators that generate phase-related linear effects, gradually damping oscillations (Fig. 6A). This framework can help refine Waddington’s epigenetic landscape and its modern iterations [48] by offering a new mechanistic metaphor of the developmental chreods. There are no rolling marbles anymore; instead, the landscape itself takes centre stage, with hills and cavities representing chromatin structure and varying densities of epigenetic marks (Fig. 6B; Additional file 2: Vid. S3). Beneath this dynamic surface, instead of Waddington’s ropes of ‘chemical forces’, multiple interconnected ratchets—representing molecular oscillators—continuously add or remove epigenetic marks. These networks enable simultaneous and highly coordinated changes across multiple loci, reshaping the landscape into new hills and valleys. Refined over millennia, this mechanism may offer a simple yet robust means of orchestrating directional, synchronized, and pre-determined cell molecular transitions during differentiation.

**Fig. 6.**
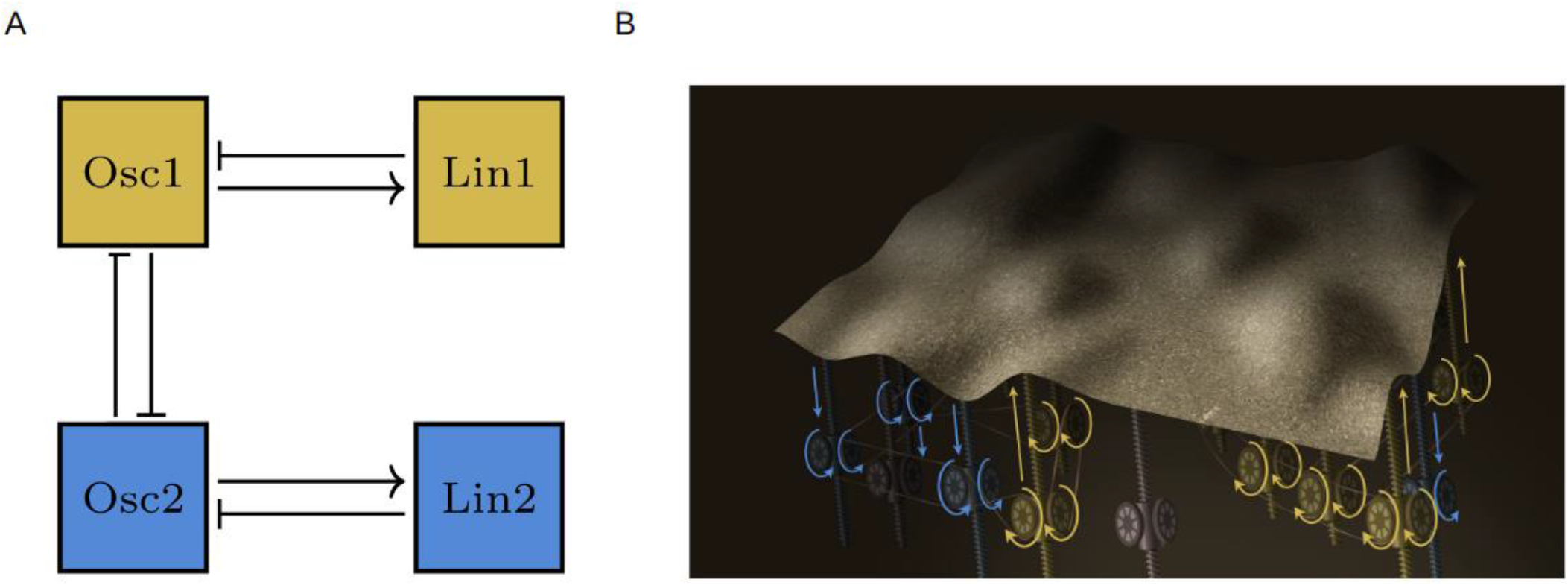
The formal and metaphorical CHREOD. **A** A repulsive interaction of the two excitable communities of oscillators, Osc 1 and Osc 2, produces antiphasic oscillations. Oscillatory cycles induce opposing linear effects, Lin 1 and Lin 2. Linear effects gradually degrade the oscillators, and the system eventually settles into a new steady-state. **B** Interconnected ratchets moving clockwise and counterclockwise represent phasic and antiphasic oscillations that alter the epigenomic landscape of developing cells in a synchronized and coordinated manner.

## Conclusions

Our studies in mouse intestinal organoids and *C. elegans* indicate that epigenomic and transcriptomic trajectories in developing cells are coupled with oscillatory dynamics. Rather than progressing strictly linearly, cells may rely on oscillations to encode and interpret temporal information. We propose that such oscillatory control extends beyond DNA modifications to include RNA, transcription factors, metabolites, and other elements of the multilevel system driving differentiation. Despite these advances, several limitations need to be discussed and addressed in future studies. Neither linear nor oscillatory measurements were fully reliable for defining precise developmental states, and the stages selected may not have been optimal for isolating oscillatory–linear dynamics. We further demonstrate that single cell approaches are promising for dealing with mixed cell populations, yet technical constraints of single-cell methods, and undetected heterogeneity may have obscured signals or added complexity. Moreover, while the CHREOD framework posits that oscillations drive linear changes, strong evidence of a causal link has not yet been demonstrated. Establishing causality will require experiments that monitor oscillations across cell cycle phases, assess interactions among competing regulatory systems, and test how perturbing specific oscillators influences both linear trajectories and the broader oscillatory network.

## Methods

### Generation and collection of mouse intestinal organoids

C57BL/6J (Strain #000664-JAX) male mice (postnatal day ∼35) were entrained to a 24 hour light–dark cycle maintained at 12 h light (7 AM) and 12 h dark (7 PM), where ZT 0 refers to lights on, for a minimum of 30 days. Mice were given ad libitum access to food, water, and a 17 cm diameter running wheel mounted in the cage. Following cervical dislocation, the abdomen of the mouse was disinfected with 70% ethanol, and an incision was made into the abdominal cavity cranial to the external genitalia, to extract the small intestine from duodenum to the pyloric sphincter. The first third of the small intestine (duodenum) were used to generate intestinal organoids. All animal procedures were approved by the Institutional Animal Care Committee of CAMH and compiled per the requirements of the Canadian Council on Animal Care and Province of Ontario Animals for Research Act.

The small intestine was opened lengthwise and cleaned in ice-cold PBS + 1X PenStrep. The tissue was chopped with a razor into ∼0.2 cm pieces, placed in a 50 mL conical tube with 30 mL of cold EDTA-shaking buffer (PBS + 15 mM EDTA + 1X Pen-Strep), and incubated for 30 minutes on an orbital shaker. The tube was then vortexed at maximum speed for 30 seconds and rested on ice for 30 seconds to detach the intestinal crypts. This was repeated 4 times. The extracted crypts were then filtered through 100 μm and 70 μm filters, and visualized and counted. The crypts were resuspended in 2:1 Matrigel to Complete Minigut Media solution (CMM; advanced DMEM/F12 supplemented with 500 ng/mL R-spondin1; 100 ng/mL Noggin; 100 ng/mL epidermal growth factor [EGF]) to a concentration of 150 crypts per 25 μL of Matrigel:CMM. CMM was supplemented with 10 μM Y-127632 ROCK inhibitor to assist with the survival of intestinal stem cells. Approximately 150 crypts were seeded into each cell culture well, incubated for 15 minutes for Matrigel to solidify, and then covered with sufficient CMM + 10 μM Y-127632 ROCK inhibitor.

In the days leading to circadian collection, the media was changed on day 1 (ZT12), day 3 (ZT6), and day 5 (ZT6) with Complete Minigut Media. Organoids were then allowed to recover for 6 hours prior to the first collection time point on day 5 (ZT12). Three to four culture wells containing 250-400 organoids (approx. 350,000 cells) were collected as biological replicates every 3-4 hours for up to 32 hours. Each sample generated up to approximately 5 μg of DNA for cytosine modification.

### Verification of circadian genes in organoids

To determine whether organoids retain circadian rhythm, we generated organoids from mPeriod1-Venus transgenic mice (MMRRC Strain #032820-JAX). Using VivaView FL incubator microscope (20X), fluorescence signals from individual organoids were measured every 20 minutes from day 2 for 6 days. mPeriod1-Venus signal intensity was used for analysis.

For qPCR, total RNA was extracted using Qiagen RNeasy Micro Kit with Qiagen RNase-free DNase I. RNA yield was quantified using NanoDrop ND-1000 (Thermo Fisher Scientific), and RNA integrity was verified via the Agilent Bioanalyzer 2100 system (Agilent Technologies). Purified RNA was converted to cDNA using High Capacity RNA-to-cDNA Kit (Life Technologies). *Arntl* (Mm00500226_m1), *Per1* (Mm00501813_m1), *Per2* (Mm00478099_m1), and *Villin-1* (Mm00494146_m1) mRNA levels were quantified using TaqMan Gene Expression Master Mix (Life Technologies) using Applied Biosystems ViiA 7 real-time PCR system. The enterocyte marker *Villin-1* was used as endogenous control. ΔΔCt was used to calculate the relative changes in transcript levels.

### Targeted bisulfite sequencing using padlock probes

Organoids were harvested using ice-cold PBS to dissolve the Matrigel and downstream processing performed in technical triplicate. DNA was purified using phenol-chloroform following proteinase K treatment. Isolated DNA was bisulfite converted using Zymo DNA Methylation kit with additional PCR cycles to ensure complete conversion. Targeted capture of bisulfite converted DNA was performed with custom design padlock probes (n = 88,007 targets).

Bisulfite padlock probes were designed using ppDesigner 2.0 [19] on the mouse reference genome (mm10) after masking for genomic variations and repeats (dbSNP 138, Microsatellites, RepeatMasker, Segmental Dups, Simple Repeats, and WindowMasker + SDust). Seven bp UMIs were added to the common sequence of each probe immediately adjacent to the probe annealing arms and later used for removal of PCR duplicates. The probes were printed by CustomArray (Bothell, WA, USA) and prepared according to the published protocol [19].

Fine-mapping of DNA modification using bisulfite padlock probes was performed using a modified version of the previously described protocol [19]. In brief, genomic DNA for each sample was bisulfite-converted and purified using the EZ DNA Methylation-Lightning Kit, with the following modifications suggested by the manufacturer for a more stringent conversion: 7.5 μL of M-dilution buffer was used for the reaction, which was incubated at 42°C for 30 min prior to addition of the CT-Conversion Reagent. A total of 185 μL of the M-dilution buffer was used in the preparation of the CT-Conversion Reagent, and only 97.5 μL of the reagent was added per reaction. The 200 ng OF bisulfite-converted DNA was hybridized to the padlock probes [19]. Targeted regions were extended using PfuTurbo Cx (Agilent Technologies) and circularization was completed using Ampligase (Epicentre). Non-circularized DNA was digested using an exonuclease cocktail and the remaining circularized DNA was amplified using a common linker sequence in the padlock probe. Libraries were PCR amplified, pooled, purified by QIAquick Gel Extraction kit and quantified by quantitative PCR on a ViiA 7 Real-time PCR system. Next-generation sequencing of the libraries was performed on either Illumina HiSeq 2500 or Illumina NovaSeq 4000 on site or at the Donnelly Sequencing Centre in Toronto, Canada.

### Preprocessing of targeted bisulfite sequencing data

For each FASTQ file, the unique molecular identifiers (UMIs) were removed from the start of the reads and saved for later processing. The FASTQ reads were quality trimmed using Trimmomatic [49] for trailing bases with a phred score < 30 and all reads with post-trimming length < 50 base pairs (bp). The trimmed reads were then aligned to a masked genome (GRCm38/mm10; mapping only to regions within a 100 bp window of the known probe locations) using Bismark v0.14.3 [50] and Bowtie 2 v2.2.2 [51]. The aligned reads were further filtered to only include reads that had start and end positions matching the padlock probe annealing arm sequence with no more than one mismatch. The filtered reads were subsequently PCR de-duplicated using the UMIs. The filtered read pairs were then converted to modification calls using the Bismark methylation extractor tool [50].

Individual cytosines were required to have a minimum coverage of 30 reads in each sample for inclusion in the analysis. β-Values were calculated as the proportion of cytosines and thymines for a given CpG; β= M/(M + U), where M = number of modified cytosines and U = number of thymines. Two samples were removed due to low sequencing coverages. All samples for a given experimental batch of organoids were internally correlated and assessed with PCA to identify outliers, where samples with average inter-sample correlation value or PC1-4 value more than 3 times the standard deviation from the mean were excluded from further analysis. To ensure equal data quality across samples we removed samples with only one remaining technical replicate from downstream analysis. Modification densities of technical replicates were median averaged and used for downstream analysis.

### scRNA-seq preprocessing

The procedure for estimating each cell population’s average TPM was as described in the adult worm study [25] — UMI counts were divided by total count for each cell, then a gene-wise average was taken which was scaled by its sum and multiplied by 10^6^. Cell-wise transcript abundance of the larval scRNA-seq dataset was obtained according to the original larval scRNA-seq study [24], where UMI counts were scaled by their size factors and log transformed and all genes with more than one UMI were analyzed.

### Cosinor analysis

A harmonic regression model was used to identify oscillations. The period (τ) was fixed to 24 h for organoid analysis, and 7 h for *C. elegans* analysis, where the phase and amplitude were modeled as a linear combination of sine and cosine terms as follows:

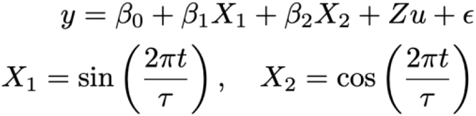

where *y* is the observed modification or normalized transcript abundance, β_0−2_ are regression coefficients, *t* is the time of observation, *Z* are the included covariates and their coefficients (*u*), and *ε* is the error term. *P*-values were obtained by comparing this model to the null model without the sine and cosine terms using an F-test. For modC analysis of organoids, a covariate for each batch and their linearity (*t*) were added to mitigate confounding effects, while the average linearity was used as the estimate of developmental linear change. Correction for multiple comparisons was performed using the Benjamini-Hochberg FDR [52] procedure. Amplitude (*A*) and acrophase (*ϕ*) in units of hours were obtained as:

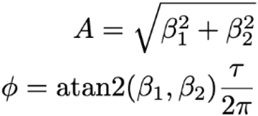

For organoid modC analysis, to determine whether the number of detected oscillating modCs was higher than expected by chance, 10,000 permutations were performed by shuffling the oscillatory component *X*_*1*_*/X*_*2*_ pairs across samples and the number of detected oscillating modCs was counted for each permutation. The permutation *p*-value was derived as the fraction of permutations with more detected oscillations than the observed.

Differences in oscillations between cell types was tested by using cosinor analysis and including an interaction term for cell type in the model. An F-test was performed between this and the cosinor model without an interaction term.

### Oscillatory-linear associations

A linear correlation of outcome and oscillation phase is inappropriate since acrophase is a circular variable. We implemented a simple association test that tests for phase associations by applying cosinor analysis across genes, where *t* = acrophase (*ϕ*) from the previous cosinor analysis and y = linearity such that any phase (*ϕ*^*^) may be associated with most up-regulation. The model was tested with an F-test.

For tissue-level analysis the degree of tissue specificity (i.e. the ratio between the tissue with the highest number of transcripts and the second highest tissue as obtained from the original study [24]) was included in the full model as a covariate for all association tests then reporting 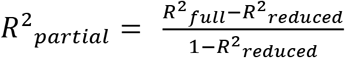 to remove the variation explained by specificity alone.

### Pause model

The pause model assumes that a continuous signal alternates oscillatory and pause states where oscillations are in circadian synchrony with the rest of the organoid. Therefore, during oscillations, the level of modC progresses as a simple harmonic oscillator with a fixed period, amplitude, and acrophase as:

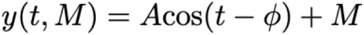

The pause start (*θ*_1_) and end (*θ*_2_) each occur at the same circadian timing, once per cycle. The oscillation after the pause is in continuity with the modC level before the pause such that:

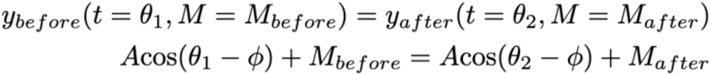

Therefore, the simplified resulting change in MESOR, in %modC/day is given in terms of a pause centre (*θ*_*mid*_) and duration (*Δθ*) by:

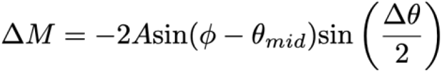

This rate of change (*ΔM*) was correlated to the observed average linearity in the cosinor analysis of modCs to obtain correlation and MSE values for each tested pause window.

### Cycle reconstructions

The cells were projected onto the first two principal components of their cell type using a truncated singular value decomposition [53,54] on the centred expression matrix and an ellipse was fit to this 2d distribution. A change of coordinates was performed such that the new coordinates (*x,y*) were given by

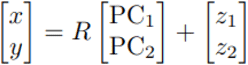

in which the rotation matrix *R* and centering vector (*z*_1_,*z*_2_) were chosen so that an unrotated ellipse could be fit in the new (*x,y*) coordinates. These parameters were computed for each cell type by first solving an ordinary least squares problem

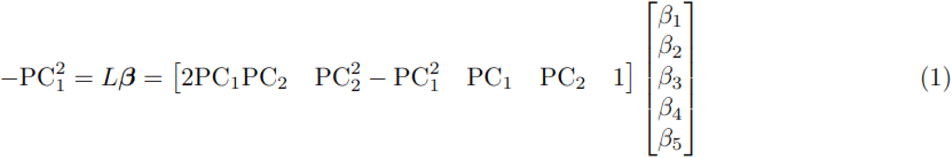

and then using the estimates 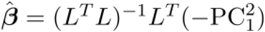 to obtain the parameters (*R,z*_1_,*z*_2_)as follows

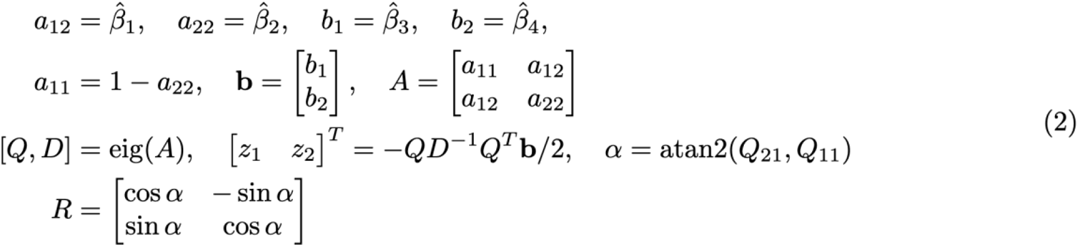

Equations (1-2) are an example of algebraic ellipse regression, and a derivation of this method can be found in ref [55]. Working in these coordinates, an unrotated ellipse was fit to the cells using the following linear regression equation

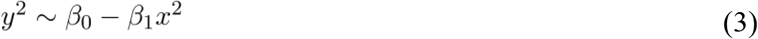

under the constraints *β*_0_ > 0 and *β*_1_ > 0, i.e. as a non-negative least squares fit. The parameters and in this model were estimated using the Lawson-Hanson algorithm [56,57] and goodness of fit was quantified using the R^2^ of the linear model in Eq. (3).

To determine whether the variance explained by ellipses was higher than expected by chance, 10,000 permutations were performed by shuffling (PC_1_,PC_2_) pairs and obtaining an ellipse fit R^2^ in each permutation. The permutation *p*-value was derived as a fraction of permutations with the permuted mean R^2^ value greater than the observed. The phase of each cell in the oscillatory cell types was estimated by finding the nearest point on the ellipse using Brent’s algorithm [58,59] and calculating the angle between the predicted point and the semi-major axis of the ellipse 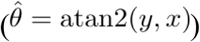. Phases were assumed to correspond to a 7 h periodicity and converted from radians to 7 h units. All genes with observed counts were tested for oscillations by cosinor analysis, using the estimated phase of the cells as *t*.

### Phase alignment (reflection and rotation)

To determine the rotation (e.g. inferred phase 1 h may correspond to observed time 6 h) and/or reflection (e.g. inferred phases 1, 2, 3 h may correspond to 3, 2, 1 h, respectively) required to align the inferred phases with the whole-worm results, we minimized the mean absolute minimum arc length (MAMAL) across genes oscillating between the whole-worm and each individual cell type. First, all minimum arc lengths — the shortest angular distance along the circle between the two phases — were obtained, where the circular mean of the arc lengths gave the phase shift required to minimize the MAMAL. This minimization was performed twice: once assuming a clockwise progression around the ellipse, and the other, a counterclockwise progression (time reversal). The direction with the smaller MAMAL was taken as the phase shift and direction required to obtain the aligned acrophase estimates.

## Data and materials availability

scRNA-seq datasets were obtained from published archives for L2 (GSM4318946) and adult (https://doi.org/10.5281/zenodo.7296546) *C. elegans*. Whole-worm larval oscillation parameters were obtained by applying the methods of the original publication to the corresponding dataset (GSE130782) and cross referenced to Dataset EV1 of the study [23]. Intestinal organoid epigenomic data will be submitted to GEO upon publication. If any additional information is required to reanalyze the data reported in this paper, contact the corresponding author, Dr. Art Petronis

## Declarations

### Ethics approval and consent to participate

Our study employed only C57BL/6J (Strain #000664-JAX) male mice (postnatal day ∼35) and transgenic mice (MMRRC Strain #032820-JAX). All animal procedures were approved by the Institutional Animal Care Committee of CAMH and compiled per the requirements of the Canadian Council on Animal Care and Province of Ontario Animals for Research Act.

### Consent for publication

Not applicable.

### Authors’ contributions

Conceptualization, A.P., M.C., E.S.O., and T.S. Cell and molecular experiments, E.S.O. Data analysis, bioinformatics, modeling, and integration, M.C., T.S., E.S.O., A.R.S., and A.P. Literature search and analysis, M.Ž. Writing of the manuscript, reviewing, editing: M.C., E.S.O., T.S., A.R.S., M.Ž., and A.P. Visualization, M.C., E.S.O, T.S., M.Ž., and A.P. Funding acquisition, A.P. All authors read and approved the final manuscript

## Acknowledgements

We thank Karolis Koncevičius for constructive feedback and discussion on the manuscript. We thank Aiping Zhang, Mrinal Pal, Daniel Groot, and Miki Susic for technical support.

