## Additional file 1 for "The chronology of developing cells: are epigenomic and transcriptomic oscillations linked to their linear trajectories?"

**Table of contents**

Fig.S1 Examples of studies showing linear dynamics of oscillating factors.
Fig.S2 Examples of oscillating modCs with and without linearity.
Fig.S3 Linearity for organoid oscillating modCs tested for significant increases or decreases.
Fig.S4 The pause model does not change correlation with pause duration.
Fig.S5 *C. elegans* linearity estimation summary.
Fig.S6 Oscillation detection in *C. elegans* scRNA-seq single-time dataset using ellipse fits to PC1-PC2.
Fig.S7 Number of detected oscillating mRNAs (X-axis; FDR q < 0.05) for each cell type with whole-worm oscillations as red and new oscillations as grey.
Fig.S8 Cell level oscillatory-linear associations and acrophase correlations prior to acrophase adjustments.
Fig.S9 Differential rhythmicity between cell types of *C. elegans* larvae.

Fig.S1

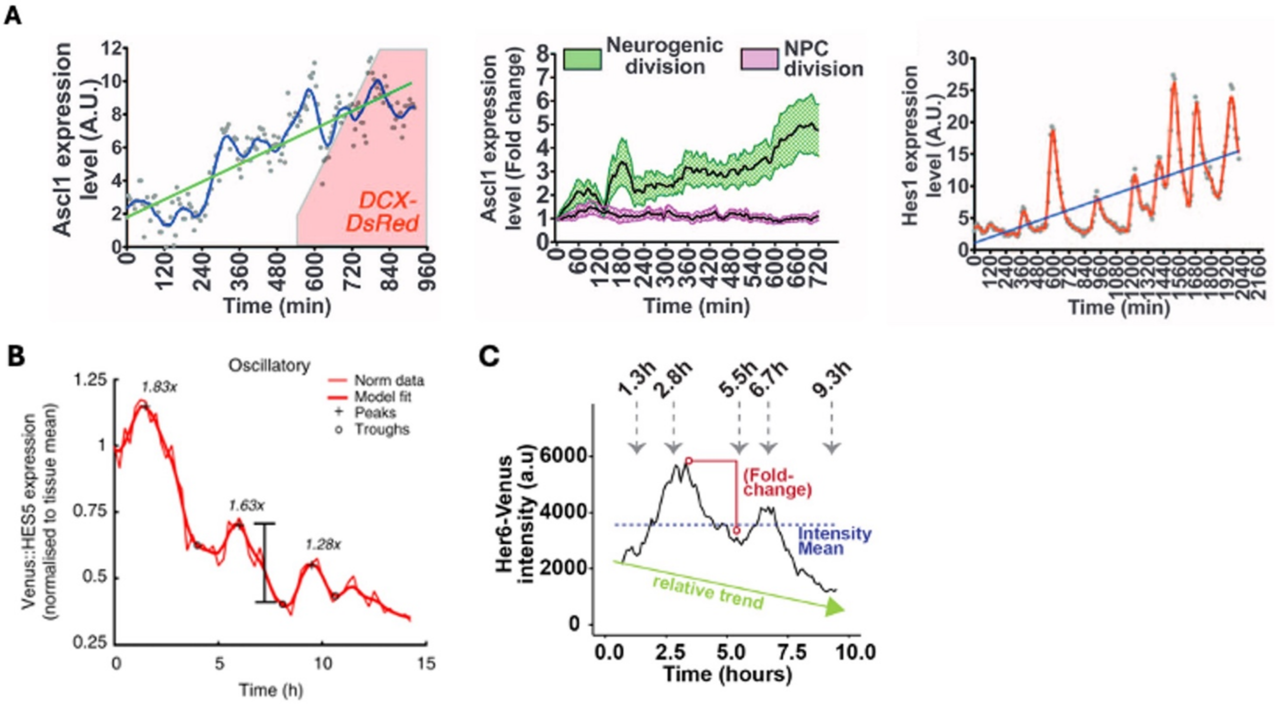


**Fig.S1 Examples of studies showing linear dynamics of oscillating factors.**

**A.** Fig. 3A, 3B and 4B of Imayoshi et al. [1] The study found sustained Ascl1 expression in differentiating neurons by using a Luc2-Ascl1 construct in mouse neural progenitors (NPCs). 6 to 8 h into the experiment, the early neuronal marker doublecortin (DCX) was expressed. In the same study, Imayoshi et al. provided evidence for Hes1 oscillations with a linear trend (Fig. 4B) during astrocyte differentiation in NPCs. The gradual development of the astrocyte identity was confirmed by tracking the expression of the astrocyte marker glial fibrillary acidic protein (GFAP).

**B.** Fig. 7B of Manning et al. [2] which identified a higher incidence of oscillatory Venus::HES5 NPC cells in mouse ventral spinal cord. As cells transitioned from NPCs to interneurons, they obtained a declining oscillatory HES5 expression profile.

**C.** Fig. 2D of Doostdar et al. [3] which identified Her6 as an oscillating bHLH transcription factor with a relative oscillatory downward trend in developing zebrafish telencephalon.

Fig.S2

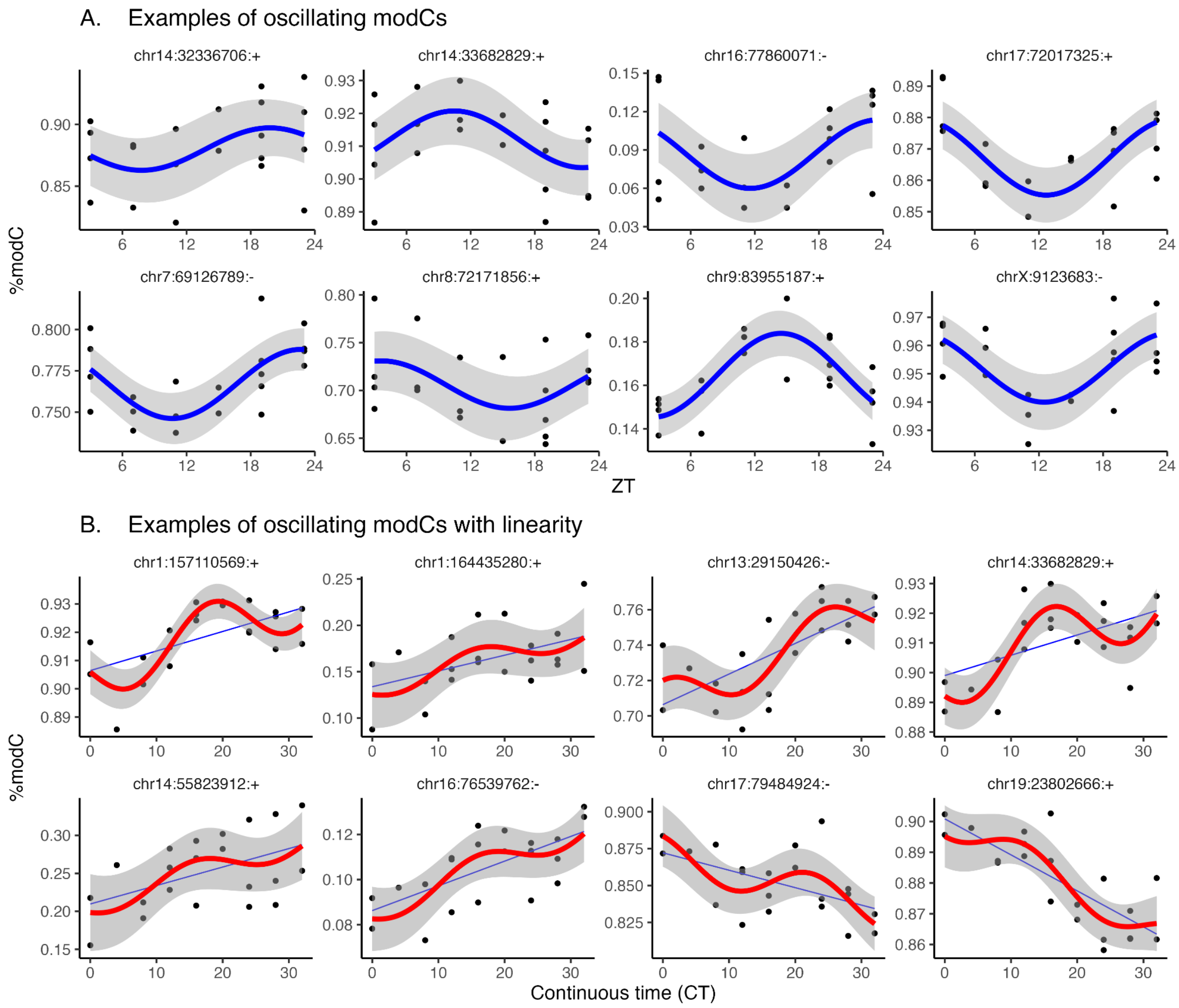


**Fig.S2 Examples of oscillating modCs with and without linearity.**

**A.** Top 8 significant oscillating modCs based on cosinor *p*-values were selected for visualization. Blue curves are cosinor fits with 95% confidence bands.

**B.** Top 8 linearly changing modCs based on linear regression *p*-value (see Additional file 1: Fig. S3) that were also significantly oscillating (cosinor $p < 0.05$) were selected for visualization. Red curves are cosinor fits with a linear covariate (time in culture) with 95% confidence bands and blue lines are linear regression fits.

Fig.S3

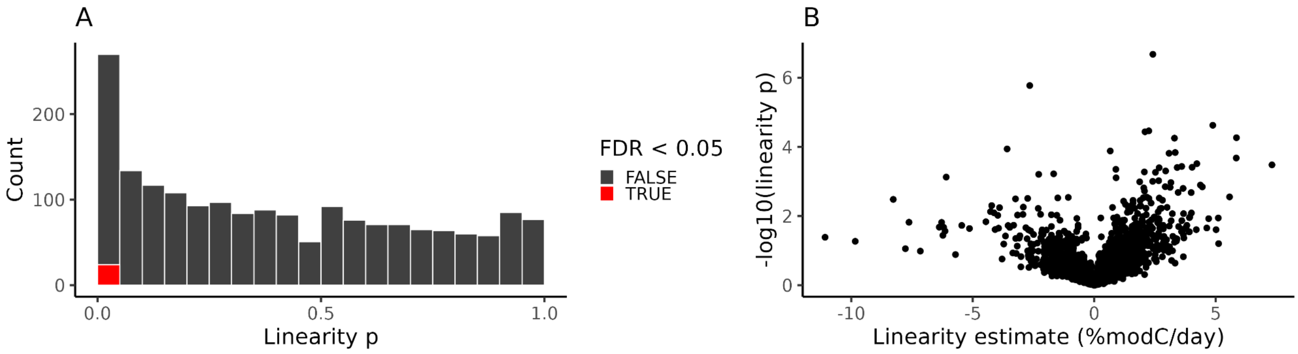


**Fig.S3 Linearity for organoid oscillating modCs tested for significant increases or decreases.**

**A, B.** Linear regression on culture time was used to demonstrate reliability of their estimates. Oscillations and batch effects were modeled as covariates, and linearity tested with an F-test with oscillations and batch effects only as the null model.

**A.** Linearity $p$-value histogram with FDR $q < 0.05$ transcripts (red).

**B.** Volcano plot of linearity estimates.

Fig.S4

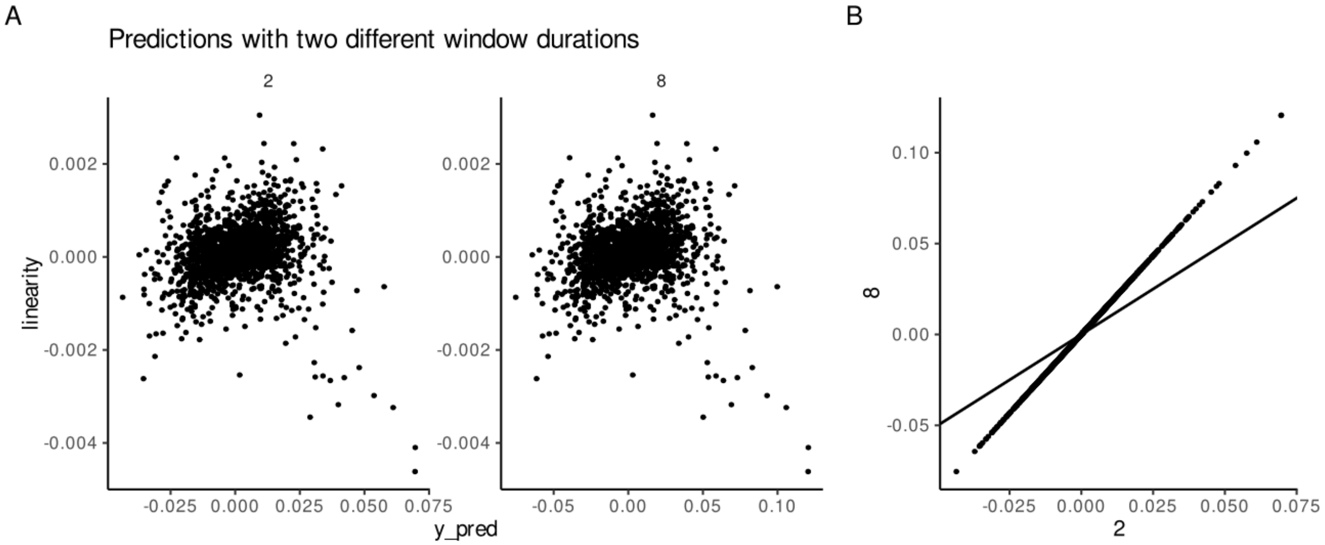


**Fig.S4 The pause model does not change correlation with pause duration.**

**A.** Correlations of model predictions (Y-axis) with observed linearity (X-axis) for two pause durations with the same window centre. Left - 2h pause, right - 8h pause. The correlations (r) are equal, but the predicted magnitudes differ.

**B.** Directly showing that predictions of two pause durations are directly proportional to one another with only a difference in magnitude; solid line indicates y = x.

Fig.S5

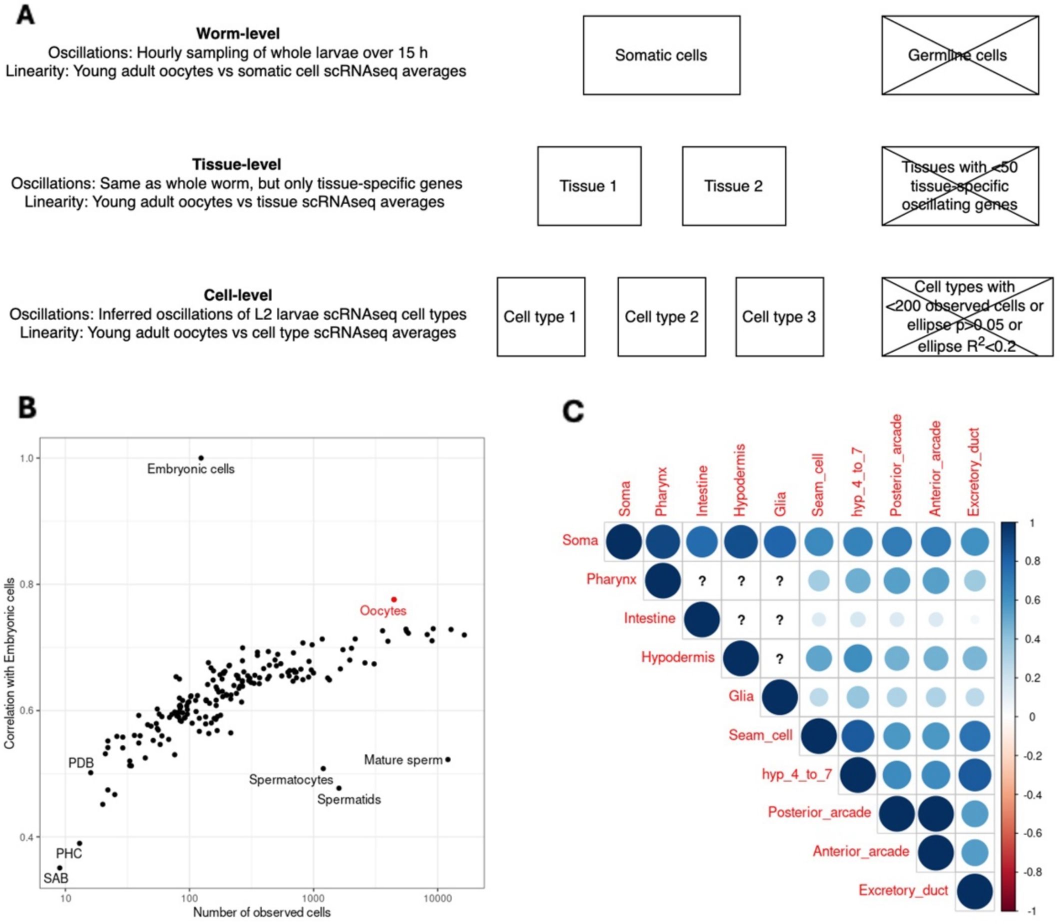


**Fig.S5 *C. elegans* linearity estimation summary.**

**A.** Left - summary of the origin of oscillatory and linearity estimates for each level of analysis. Right - The main tissue or cell exclusion criteria for each level.

**B.** Correlation with embryonic cells across all transcripts (Y-axis) versus the number of cells for the cell type on a logarithmic base 10 scale (X-axis). Note that embryonic cells were n = 124, while oocytes were n = 4,447. Oocytes were the most similar cell type to the few embryonic cells in this dataset.

**C.** Pearson correlation of linearity estimates across the 3 levels of analysis for all genes with estimates between pairwise comparisons. Young adult somatic cells were all cells not labelled as “Germline”.

Tissue-level linearity was estimated for L2 tissue labels across the following adult cell type labels: Pharynx=’Pharynx’ and ‘Arcade cells’, Intestine=Intestine, Hypodermis=’Hypodermis’ and ‘Seam’ , ‘Glia’=’Excretory’ and ‘Support cells’. Cell level linearity for each L2 cell type was calculated between the following cell types and oocytes for ‘Seam_cell’, ‘hyp_4_to_7’, ‘Posterior_arcade’, ‘Anterior_arcade’, and ‘Excretory_duct’, respectively: ‘Seam cells (bus+)’, ‘hyp7 (hypodermis)’, ‘Arcade cells’, ‘Arcade cells’, ‘Excretory duct’.

Fig.S6

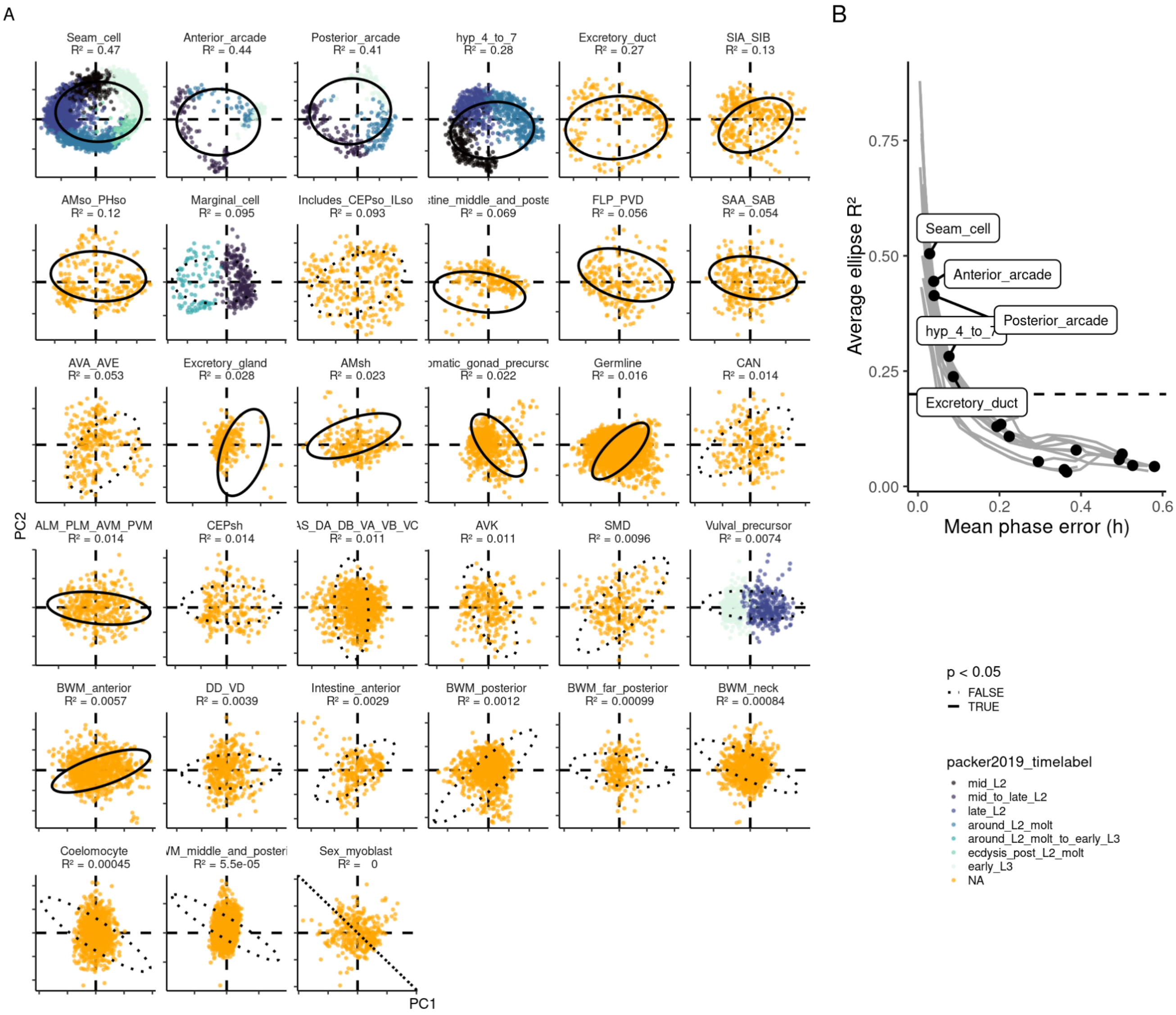


**Fig.S6 Oscillation detection in *C. elegans* scRNA-seq single-time dataset using ellipse fits to PC1-PC2.**

**A.** The cell distribution on the PC1-PC2 projection (X-/Y-axis) for ellipse regression is shown for each cell (dots). The black oval curve in each panel represents the ellipse of best fit. Permutation $p < 0.05$ ellipses are shown as solid lines with permutation $p > 0.05$ as dotted lines and panels are ordered by ellipse regression $R^2$. The color represents temporal annotations present in the original dataset for select cell types [4].

**B.** The phase reconstruction error in each cell-type was estimated by finding the mean phase error (X-axis) that corresponded to the observed $R^2$ (black dot). Mean phase error was calculated as the minimum mean absolute minimum arc length for all phase shifts between the real and estimated phase. Simulated $R^2$ values were obtained from simulated datasets with the same sample size and x-y noise ratio as the real data, i.e. for observed noise $(\sigma_0^x,\sigma_0^y)$, simulated noise was of the form $(\sigma^x,\sigma^y) = k(\sigma_0^x,\sigma_0^y)$ with scaling factor $k > 0$. The average of 20 simulations was taken for each cell type and each noise scaling factor to obtain lines estimating phase error.

Fig.S7

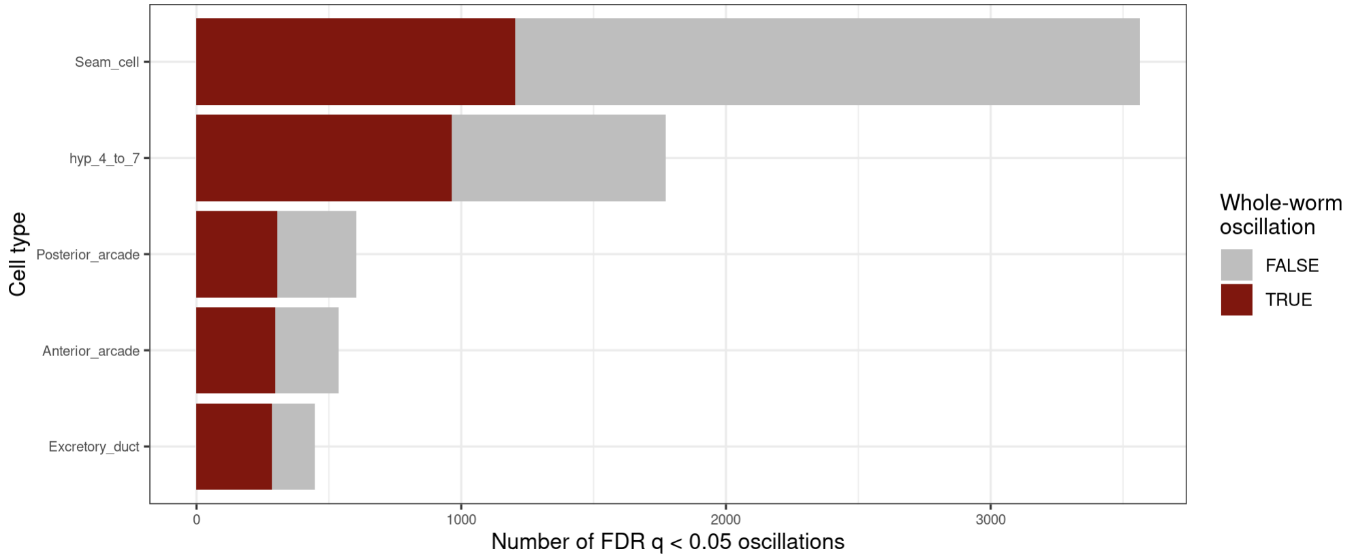


**Fig.S7 Number of detected oscillating mRNAs (X-axis; FDR *q* < 0.05) for each cell type with whole-worm oscillations as red and new oscillations as grey.**

Fig.S8

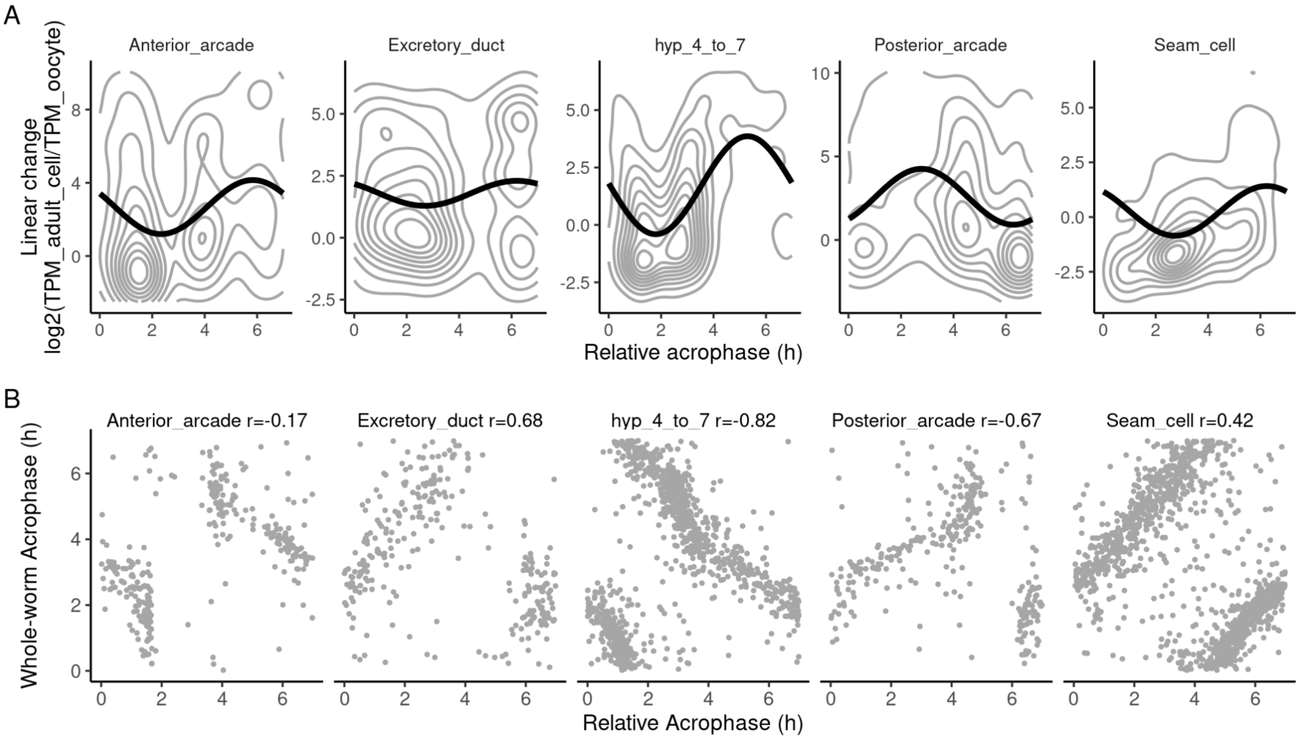


**Fig.S8 Cell level oscillatory-linear associations and acrophase correlations prior to acrophase adjustments.**

- **A.** Cell-level association between unadjusted/relative acrophases of oscillating mRNAs with a 7 h period (X-axis) and estimates of transcriptomic linear change of the corresponding adult cell type (Y-axis). Linearity was the difference in logarithm (base 2) transcripts per million between oocytes and the adult cell population. Contours (grey) were rendered using 2D kernel density estimation. The cosinor curves (black) were estimated by regression, using the sine and cosine of the acrophase as predictors of linearity. Association statistics were as seen in Fig. 5.
- **B.** Unadjusted/relative acrophases (X-axis) correlated with whole-worm acrophases (Y-axis). Circular correlation values (r) are shown in each panel which are identical to those seen in Fig. 5.

Fig.S9

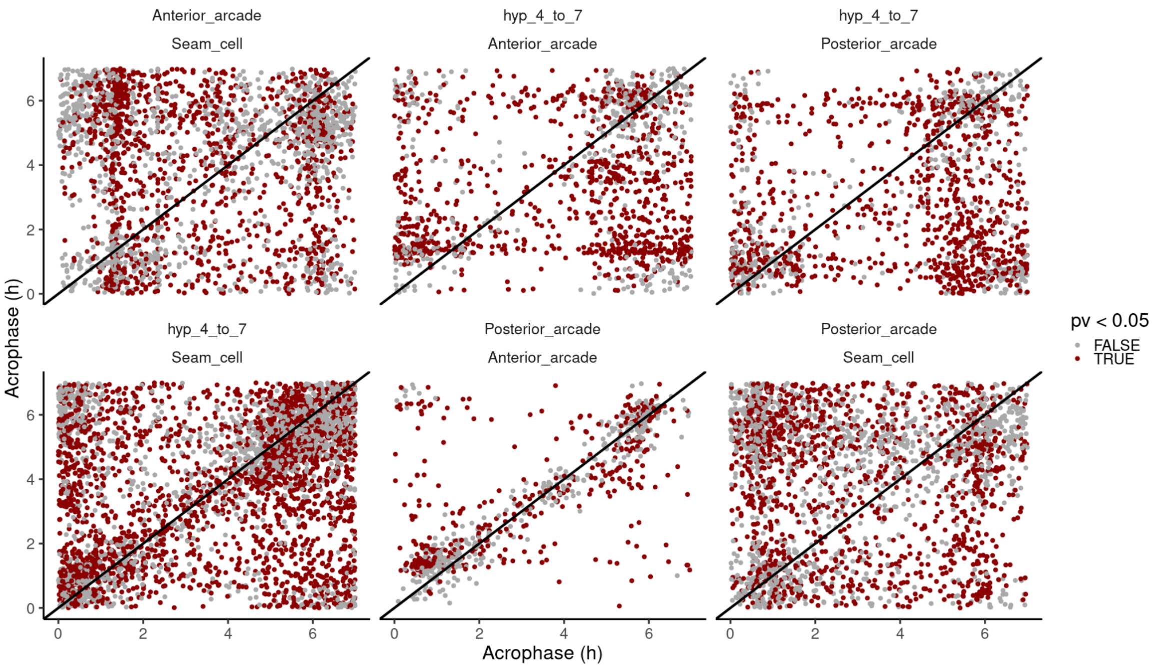


**Fig.S9 Differential rhythmicity between cell types of *C. elegans* larvae.**

Acrophase differences between pairwise cell types for genes that were oscillating in both cell types used for the calculation of average phase shift between cell types. Red dots were significant for differential oscillations at $p < 0.05$.
